# Discovering Immunoreceptor Coupling and Organization Motifs (ICOMs)

**DOI:** 10.1101/2023.07.21.550023

**Authors:** Michael Reth

## Abstract

The recently determined cryo-EM structures of the T cell antigen receptor (TCR) and B cell antigen receptor (BCR) show in molecular details the interactions of the ligand-binding part with the signaling subunits but they do not reveal the signaling mechanism of these antigen receptors. Without knowing the molecular basis of antigen sensing by these receptors, a rational design of optimal vaccines is not possible. The existence of conserved amino acids (AAs) that are not involved in the subunit interaction suggests that antigen receptors form higher complexes and/or have lateral interactors that control their activity. Here, I describe evolutionary conserved leucine zipper (LZ) motifs within the transmembrane domains (TMD) of antigen and coreceptor components that are likely to be involved in the oligomerization and lateral interaction of antigen receptor complexes on T and B cells. These immunoreceptor coupling and organization motifs (ICOMs) are also found within the TMDs of other important receptor types and viral envelope proteins. This discovery suggests that antigen receptors do not function as isolated entities but rather as part of an ICOM-based interactome that controls their nanoscale organization on resting cells and their dynamic remodeling on activated lymphocytes.

## Introduction

Over the past three years the human population has experienced a major viral pandemic with the accompanying loss of life and economic decline. What will hopefully now end this health crisis is the establishment of an adaptive T and B cell immunity against the pandemic SARS-CoV-2 virus and its mutants in vaccinated and/or infected individuals. The antigen receptors on T and B cells play an essential role in adaptive immunity. The T-cell antigen receptor (TCR) and the B-cell antigen receptor (BCR) have a similar molecular design. They are multi-protein complexes consisting of a ligand-binding part and a signaling subunit. For the αβTCR, these are the α/β chains binding to peptide-loaded MHC (pMHC) molecules and the CD3/zeta signaling complex consisting of CD3γ/CD3ε and CD3δ/CD3ε heterodimers and a TCR-ζ/TCR-ζ homodimer (1).

For the BCR, the ligand-binding part consists of the different classes of the membrane-bound immunoglobulin (mIg) and the Igα/Igβ (CD79a/CD79b) heterodimer (2). All signaling subunits of these receptors carry a dual tyrosine motif in their cytoplasmic tail, which I first described in 1989 (3) and, which became known as the immunoreceptor tyrosine-based activation motif (ITAM). It has been shown that the two ITAM tyrosines, once phosphorylated, are engaged by the dual SH2 domains of ZAP-70 and Syk on the TCR and BCR, respectively (4, 5). How ITAM phosphorylation is prevented in resting T and B cells and how exactly antigen binding to the receptors results in increased accessibility of the ITAM tyrosines for phosphorylation and kinase binding is not known at present.

The nanoscale organization of the antigen receptors on the plasma membrane seems to be essential for their regulation (6-8). Antigen-dependent activation of B cells results in a nanoscale reorganization of antigen receptors and their membrane environment. As proposed by the dissociation activation model (DAM), the IgM-BCR oligomers are opened and gain access to the coreceptor CD19 and to CD20 (9, 10). How nanoscale receptor clusters are established and what molecular events are involved in their remodeling are currently unknown. I suggest here that not only the extracellular or intracellular parts of immunoreceptors, but also their TMD play an important role in their proper function. This suggestion is supported by the discovery of sequence motifs within these TMDs that are likely to regulate the nanoscale organization of these receptors on resting and their dynamic reorganization on activated lymphocytes.

## Results

### The Asymmetric Organization of the BCR and TCR Complexes

In most immunology textbooks, TCR- and BCR complexes are depicted as symmetric structures with a signaling heterodimer placed on each side of the ligand-binding parts. Based on the fact that the mIgM molecule is a symmetric homodimer, we initially also proposed a symmetric model for the IgM-class BCR (IgM-BCR) complex, in which the mIgM molecule binds two CD79a/CD79b heterodimers (11). However, biochemical analysis has shown that the mIgM molecule and the CD79a/CD79b heterodimer form a 1:1 complex rather than a 1:2 complex (12). The now resolved cryo-EM structures confirm the 1:1 model of the IgM-BCR complex and show an asymmetric assembly between the ligand-binding and signaling part of these antigen receptors (13-15). Specifically, in the murine IgM-BCR structure the CD79a/CD79b heterodimer is predominantly interacts with only one of the two heavy chains (μHC and μHC’) of the mIgM molecule (15). The monomeric αβTCR also shows an asymmetric organization with the CD3 complex occupying only one side of the αβTCR (16). Interestingly, the cryo-EM structures of the pMHC-bound or free αβTCR do not reveal major alterations upon ligand binding (17). Coupling between the antigen receptor components occurs via three distinct contact sites, namely the membrane proximal Ig domains, the connecting peptides and the TMDs, with the latter two playing a dominant role in complex formation. The TMDs of all antigen receptor components are single-spanning alpha-helixes, covering a space within the membrane that has been previously underestimated and drawn much too small in many schematic representations of antigen receptor organization. Thus, only the recent cryo-EM structures highlight the importance of the TMDs for the formation and stability of antigen receptor complexes.

### Analysis of the Lateral Accessibility of the Monomeric IgM-BCR Complex

A major question arising from the known cryo-EM structures is why the basic structure of the antigen receptor complexes is asymmetric. A possible answer to this question is that the asymmetric organization allows these receptors to interact on the lymphocyte membrane either with themselves (forming dimers and oligomers) or with coreceptor modules that regulate or amplify the signal transduction of the antigen receptors. Following this idea, I took a closer look at the conserved TMD AAs that are either engaged in the formation of the 4-alpha-helical bundle of the IgM-BCR TMDs or exposed to the lipid bilayer. The TMDs of the CD79a and CD79b signaling component have different positions in the IgM-BCR structure. The CD79a TMD binds on one side of the alpha helix to the CD79b TMD and on the other side to the TMDs of the μHC and μHC’ (Fig.1). Indeed, it is predominantly CD79a that couples the CD79a/CD79b heterodimer to the mIgM molecule. The CD79b TMD is only engaged on one side in the binding to the CD79a TMD and, to a lesser extent, to the μHC’ (Fig. 1). The exposed, free side of the CD79b TMD carries either a leucine or isoleucine residue at the TMD position 8, 11, 15, 18, 22, which are part of a LZ heptad motif involved in the interaction of different TMD alpha-helixes (18). Within a heptad (**a**bc**d**efg) motif, the a- and d-position residues form the hydrophobic core of the interface between the interacting alpha helices.

**Figure 1.**
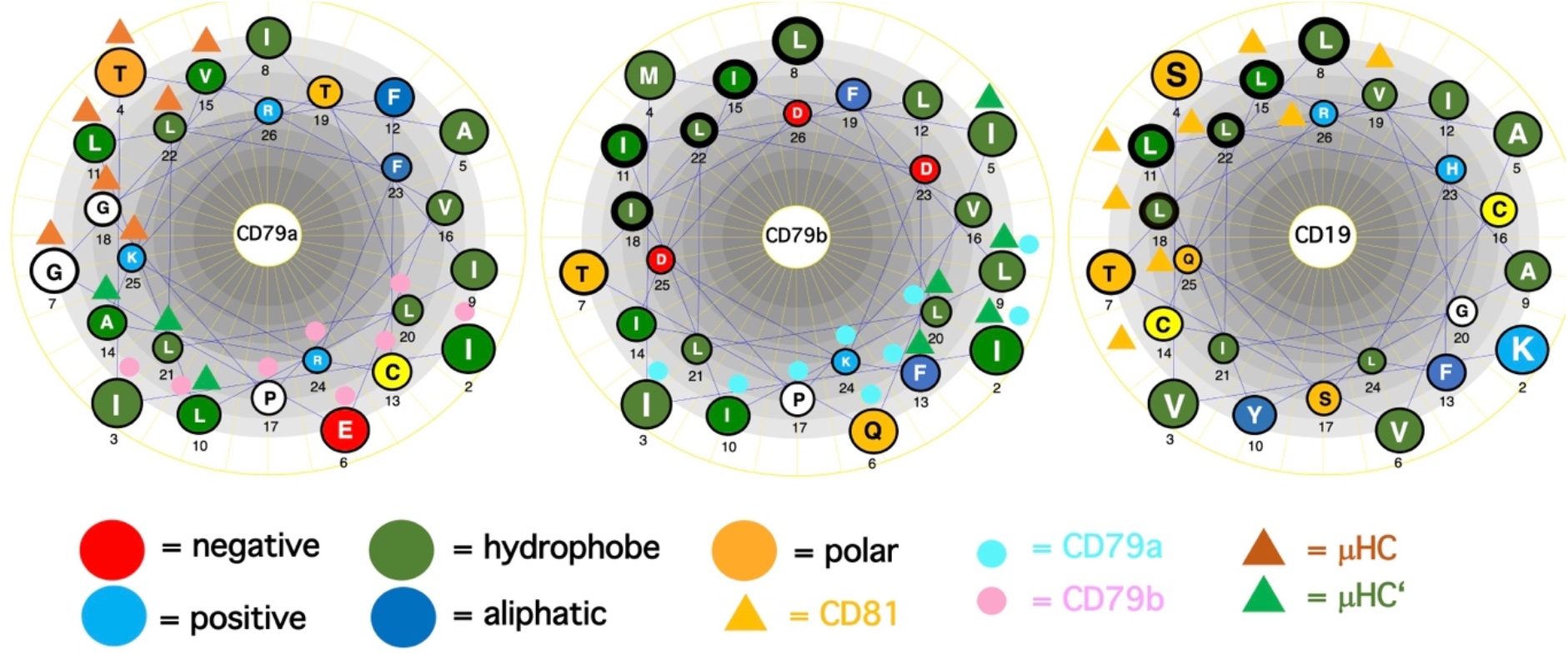
Comparison of the TMD AAs sequences (single-letter code) of the BCR signaling subunits CD79a, CD79b and of the coreceptor CD19 represented as helical wheel. The AAs are color coded according to their chemical features, as indicated. Those AAs involved in BCR or CD19/CD81 complex formation are marked by colored triangles and circles. The ICOM AAs which form the central LZ-structure are outlined by a bold circle.

### Discovery of a Potential Lateral Interaction Motif with the TMDs of Immunoreceptors

From our previous studies of the BCR conformation and interaction on the B cell membrane using the Fab-based proximity ligation assays (Fab-PLA), we learned that the IgM-BCR established a close contact with the CD19 coreceptor upon B cell activation (6). A look at the AAs composition of the human CD19 TMD revealed a series of 5 leucine residues located on one side of the CD19 TMD alpha helix at the same position as the LZ components of the CD79b TMD (Fig.1). It is thus likely that the LZ structure within the CD19 TMD is involved in the heterodimerization of the IgM-BCR with the CD19 coreceptor. Interestingly, in our Fab-PLA studies, we detected the IgM-BCR/CD19 interaction only on activated but not on resting B cells. The reason for this restriction may be the finding that CD19 is only present on the B cell surface in association with the CD81 protein (19). A recent cryo-EM study shows the molecular details of the AAs association within the CD19/CD81 complex and I found that the LZ residues of CD19 are covered by one of the TMD alpha-helixes of CD81 and are not available for binding to the CD79b TMD(20). However, B cell activation appears to be accompanied by the dissociation of the CD19/CD81 complex, releasing CD19 for the potential TMD interaction with the IgM-BCR complex (21). As explained above, the association between the ligand-binding and the signaling subunits of the IgM-BCR involves the extracellular Ig domains and the connecting peptides in addition to the TMDs. Thus, it is feasible that not only the LZ structure within the TMDs but also additional sites take part in the CD79b/CD19 complex formation. One of these sites seems to be the juxtamembrane region which contains a series of complementary positively (H23,Q25,R26) and negatively (D23,D25,D26) charged AAs in the CD19 and CD79b AA sequence, respectively (Fig.1). An alignment of the human TMD sequences of CD79b and CD19 according to the found LZ structure shows that these two sequences carry 9 identical or similar AAs (Tab.1). Thus, I could define an LZ-containing A-motif: TLxx(LI)(LI)xx(LI)xx(LI)xx(LI)L, which is shared by both proteins and that may play an important role in the organization of antigen receptors and their coupling to lateral interactors. For this and related LZ motifs (see below), I propose the name “immunoreceptor organization and coupling motifs” (ICOMs).

**Table 1:**
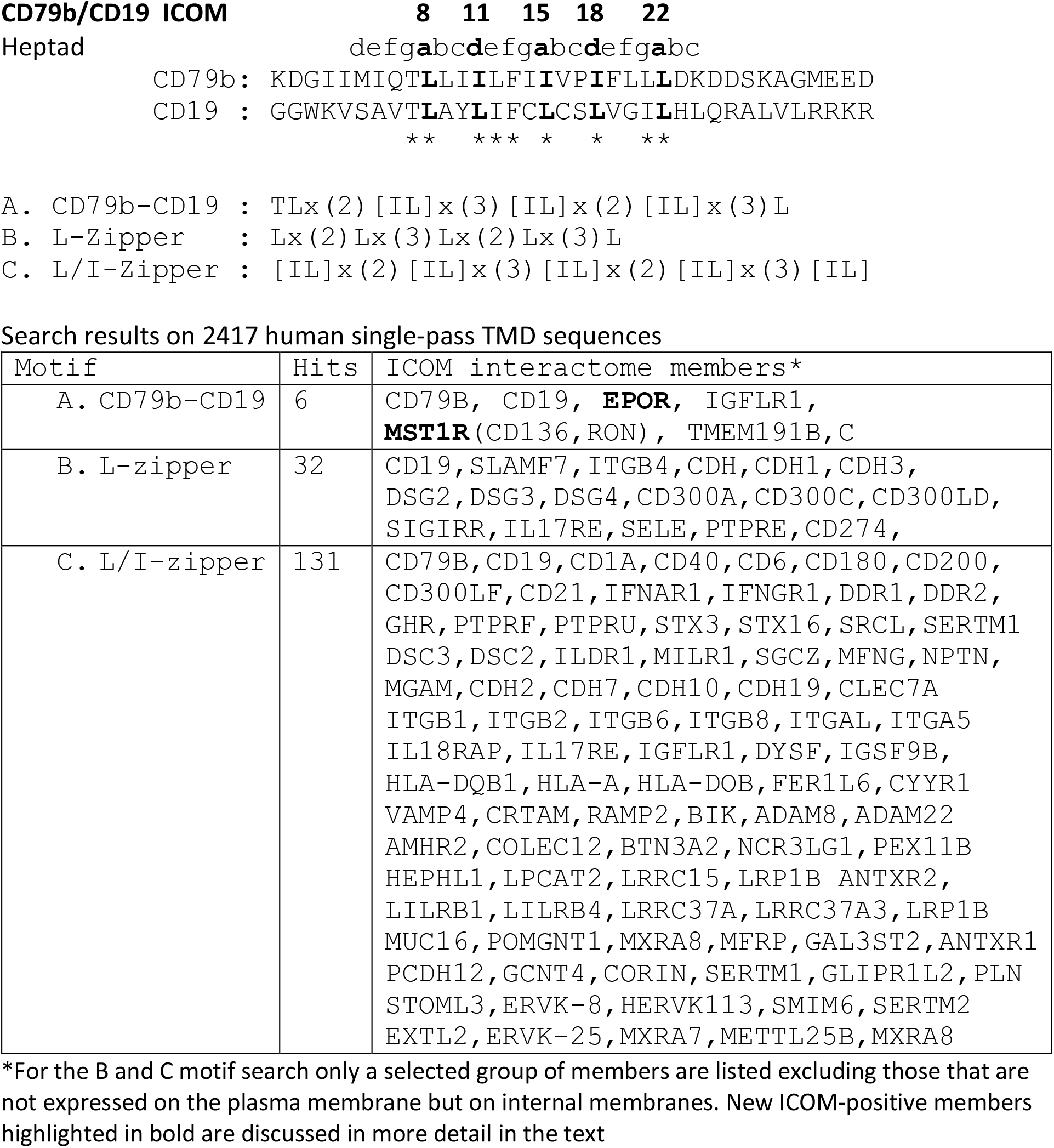
ICOMs Motif Searches related to the CD79b and CD19 TMD Sequence.

### Motif Search for ICOM-Family Members

A prosite motif search of the TMD + 5 AA flanking sequences of 2417 single-pass transmembrane proteins extracted from the human uniprot database (TMD5) for the defined A-motif yielded only 6 hits (Tab. 1). In addition to CD79b and CD19, the A-motif is found in the TMD sequence of the erythropoietin receptor (EpoR), the insulin growth factor-like receptor 1 (IGFLR1), the macrophage stimulating 1 receptor (MST1R), also known as CD136 or RON, and the transmembrane proteins TMEM191 of unknown function. The result of the A-motif search was verified by a TMD sequence comparison according to the 5 positions of the heptad LZ motif (Tab. 2). A problem that arises with such a comparison is that the start of a TMD sequence is not always well defined and that the length of this sequence can differ between 19-24 AAs. In my TMD sequence comparison, I aim to place the first heptad repeat AA at (or near) the position 8 and the last at position 22, which should be preferentially followed in the next 1-4 positions by a charged or polar AA. With such an alignment the 5 positions of the heptad repeat are occupying a central space within the alpha-helical barrel representation of a TMD sequence (Fig. 2). Of the 6 hits from the A-motif search, only the MST1R TMD caused a problem and allowed alternative ways of LZ motif sequence alignment. One should keep in mind that an LZ sequence is a rather flexible dimerization structure that is not restricted to leucine or isoleucine residues. Indeed, 1 or 2 of the 5 heptad repeat positions of an LZ structure could carry other AAs than leucine or isoleucine, such as hydrophobic or polar, without compromising the dimerization function (22-24). Cells appear to exploit this binding flexibility to either increase or decrease the affinity of an ICOM:ICOM interaction (see below).

**Table 2:**
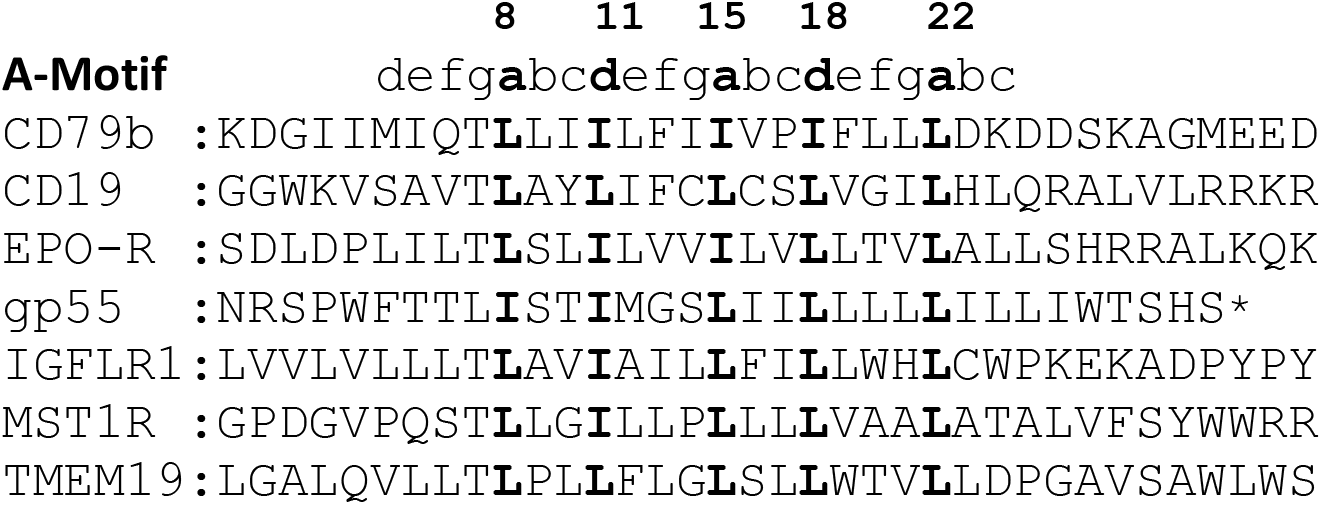
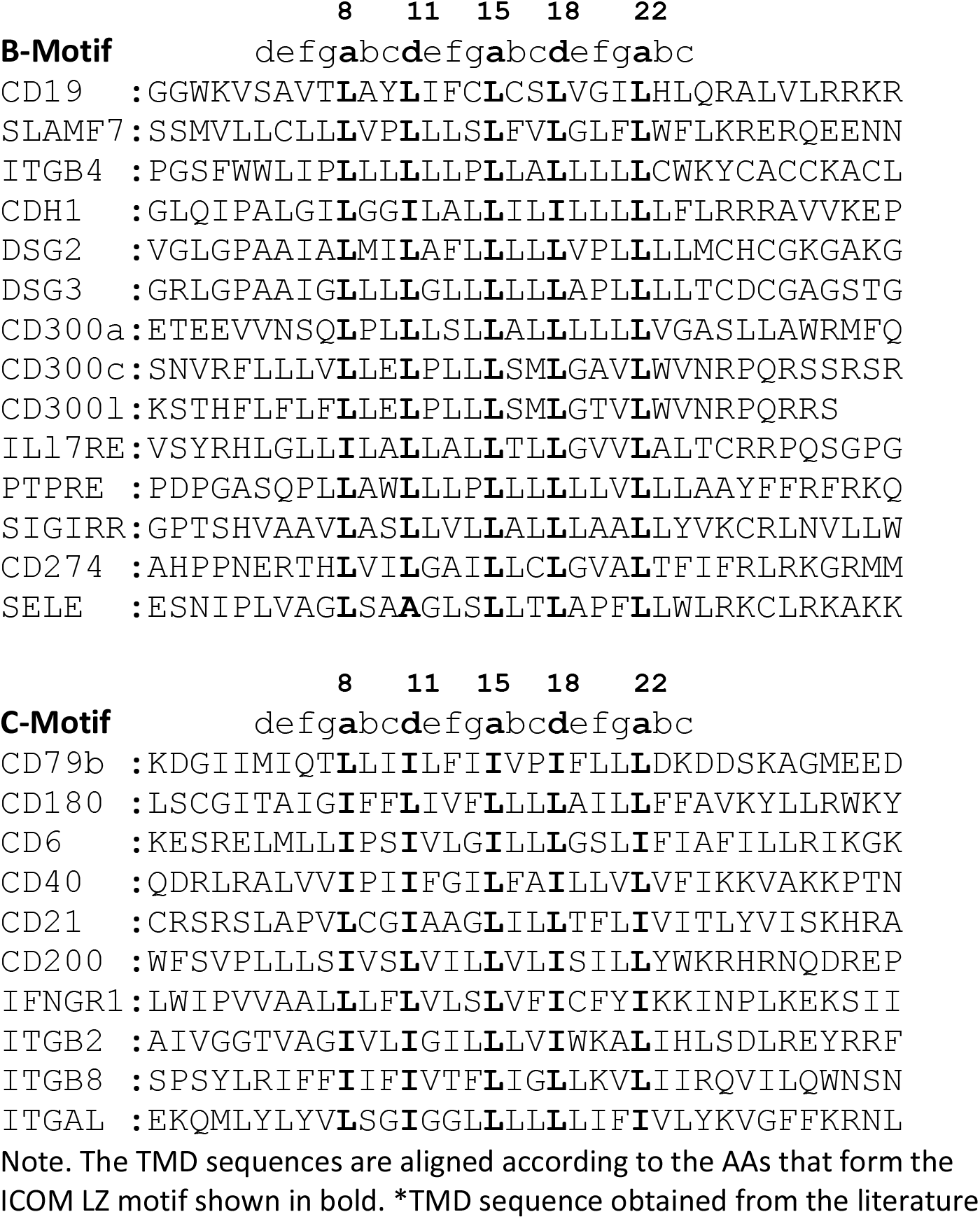
Selected TMD Sequences from the ICOM A-C Motif Searches.

**Figure 2.**
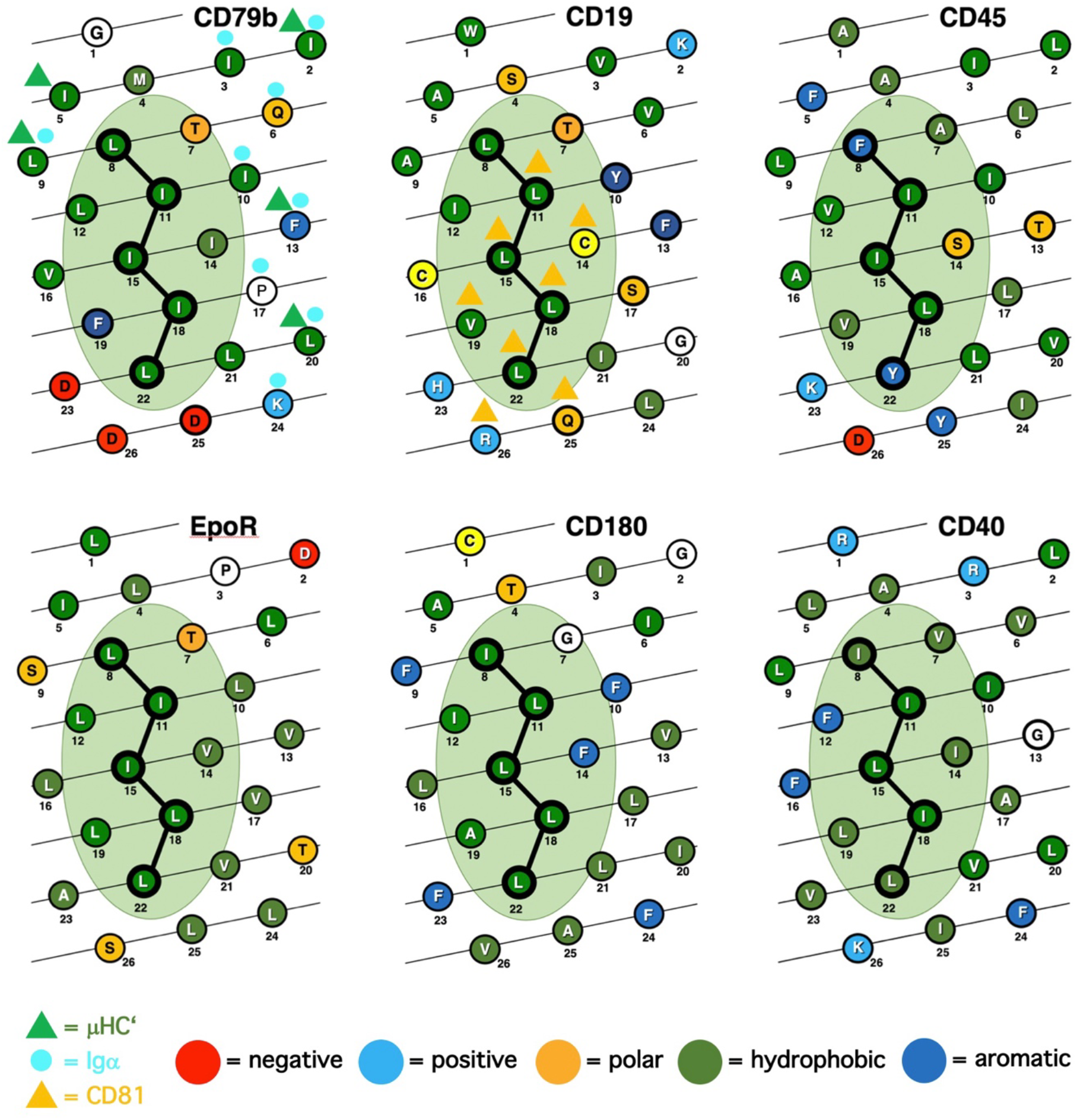
Comparison of the TMD AAs sequences (single-letter code) of ICOM-positive membrane proteins represented as alpha-helical barrel. The AAs are color coded according to their chemical features. Those AAs involved in BCR or CD19/CD81 complex formation are marked by colored triangles and circles. The ICOM AAs of the (**a**bc**d**efg) heptad motif are connected by a line and outlined by a bold circle.

Next, I conducted a prosite search for a leucine-only LZ motif (B motif): Lx(2)Lx(3)Lx(2)Lx(3)L found in the CD19 TMD. This search yielded 32 hits, including several adhesion proteins such as cadherins (CDH, CDH1, CDH3), desmogleins (DSG, DSG2, DSG4) and the integrin ITGB4. It is thus feasible that an ICOM-based interaction between adhesion and antigen receptors is involved in the activation of lymphocytes in contact with cell-bound antigens.

A prosite search for ICOM members with a mixed leucine/isoleucine LZ motif (C-motif): [IL]x(2)[IL]x(3)[IL]x(2)[IL]x(3)[IL] identified 131 membrane proteins. This search included the 32 MPs identified in the B-motif search and extended the group of adhesion proteins carrying an ICOM within their TMD. Interestingly, the identified ICOM-positive members expressed on B cells are CD40, CD180 also known as RP105 (see below). Furthermore, I identified the scavenger receptor family member CD6, which fine-tunes TCR signaling, as an ICOM-positive protein.

### The EpoR/p55/sf-Stk Story

The finding that the TMD of EpoR contains a LZ structure similar to that of CD79b and CD19 is of particular interest since this receptor has been extensively studied over the last 20 years. Like the BCR, the EpoR was once thought to be activated either by dimerization of two monomers or by an EPO-dependent conformational change that brings two TMDs into close proximity to each other (25). However, this idea has been challenged by studies showing that the resting EpoR already forms a dimer and that the LZ motif within its TMD is involved in receptor dimerization (26, 27). The EpoR can be activated not only by the binding of its ligand EPO but also by coexpression of the envelope-related glycoprotein gp55 of the Friend spleen focus forming virus (F-SFFV), which causes erythroblastosis in an EpoR-dependent manner (26) (28). Interestingly, the TMD of gp55 carries an ICOMs-related LZ motif (Tab. 2). Newer data show that the TMD of gp55 either forms homodimers or binds to the TMD of EpoR (29). Thus, dissociation of the TMDs of the EpoR dimer either by Epo or by gp55 has been discussed as the activation mechanism of this receptor. Recent studies suggest that in addition to EpoR and gp55 also a third player, namely the tyrosine kinase sf-Stk, is required for fibroblast transformation by F-SFFV by forming a tripartite EpoR/p55/sf-Stk complex as an oncogenic driver (30, 31). Interestingly, sf-Stk is an N-terminal truncated form of murine MST1R/RON, which is one of the six hits of the A-motif search of the human membrane proteome. Thus, three different ICOM-positive membrane proteins are implicated in oncogenic complex formation and cellular transformation. In summary, genetic and biochemical studies support the notion that ICOM-carrying TMDs are involved in the homo- and/or heterodimerization of membrane proteins and regulate important biological processes such as cellular transformation.

### Viral ICOMs and Their Targets

When I discovered the dual tyrosine motif (now known as ITAM) in the cytoplasmic tail of the signaling components of the BCR and TCR, as well as Fc receptors, I noticed that the glycoprotein gp30 of the bovine leukemia virus (BLV) also carried such a motif (3). Subsequently, other ITAM-bearing viral proteins were described, demonstrating that viruses employ the ITAM-dependent signaling modules as part of their propagation strategy (32).

Given the importance of the ICOM interactions for lymphocyte regulation and activation, it is likely that viruses and tumor cells employ the ICOM coupling and organization function. Indeed, I found that BLVgp30 carries not only an ITAM in its cytoplasmic tail but also an ICOM within the TMD sequence (Tab. 3). BLVgp30 may thus interact with CD19 in an ICOM-dependent manner to establish an ITAM/PI-3K signaling axis and this might be one of the drivers of bovine leukemia. Whether or not BLVgp30 or the env proteins of related viruses can also directly interact with the antigen receptors in the way that gp55 activates the EpoR remains to be investigated. Clearly, a deregulated ICOM interaction is involved not only in viral infection but also in oncogenic transformation. Another example for this is the platelet-derived growth factor receptor beta (PDGFRb), whose TMD can form ligand-independent homodimers (33) and interact with the bovine papillomavirus BPVE5R transmembrane protein (34, 35). The TMD of both proteins carries an ICOM-related LZ structure, which may promote their interaction and tumor development (Tab. 3).

**Table 3.**
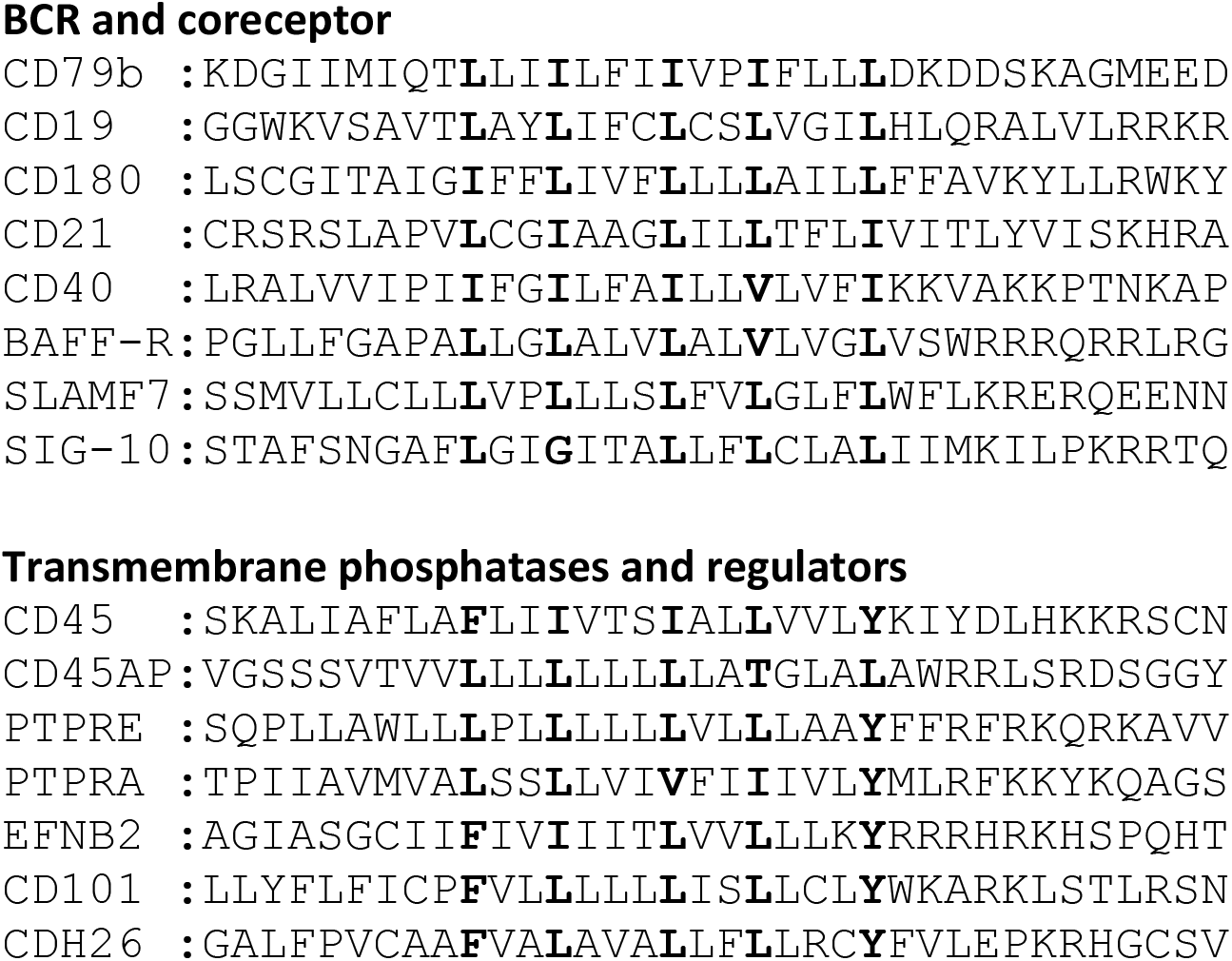
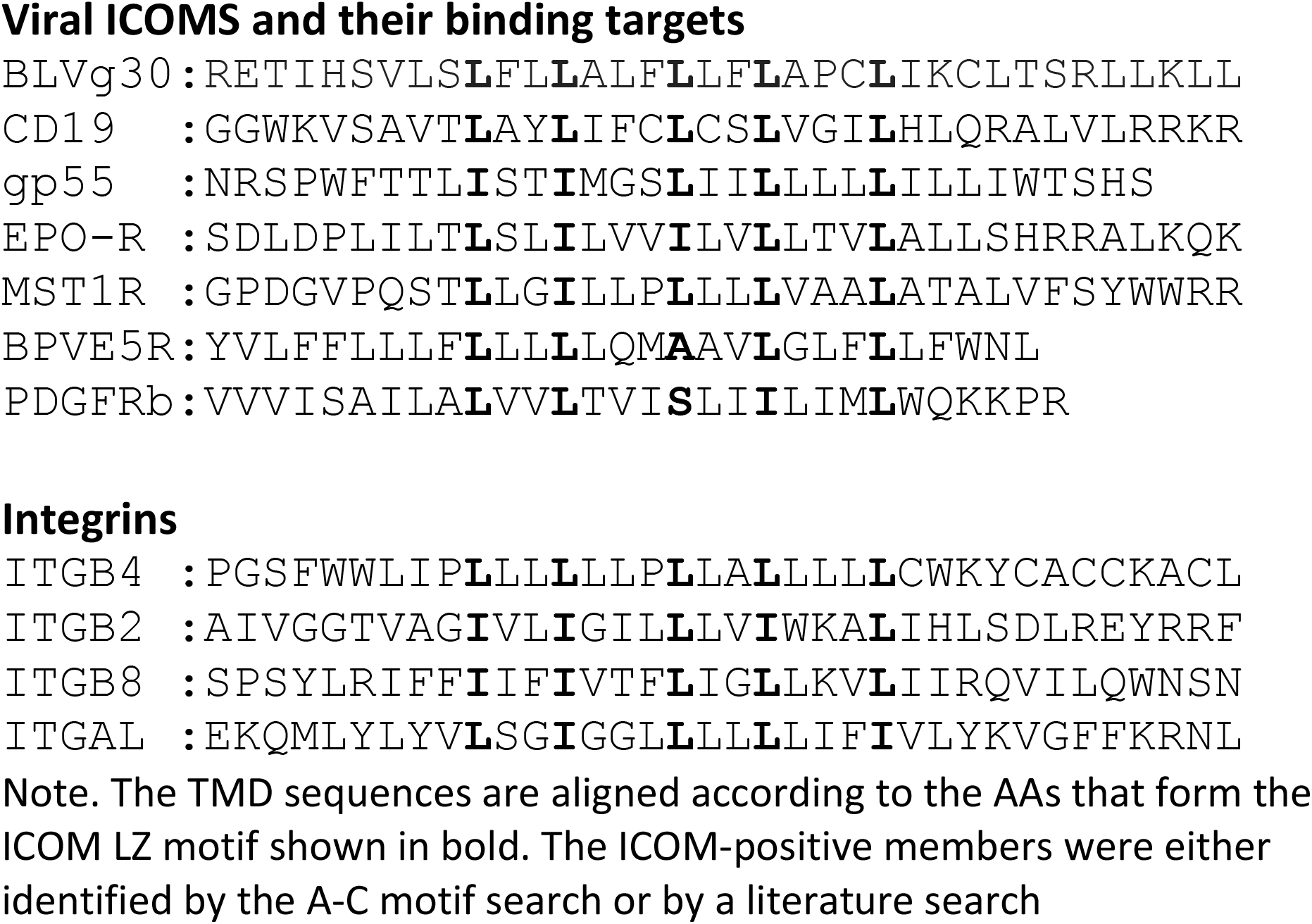
TMD Sequences of ICOM-positive Receptor Components BCR and coreceptor.

### The CD45 ICOM-Related TMD Sequence

In our nanoscale Fab-PLA studies, we found that the transmembrane phosphatase CD45 (PTPRC) is associated with the IgM-BCR complex on resting but not on active B cells. By dephosphorylating ITAM tyrosines, CD45 has a negative signaling role as a gatekeeper of IgM-BCR activation (8). However, upon B cell activation, CD45 dephosphorylates the negative regulatory pY593 of Lyn and has a positive role in IgM-BCR signal amplification. Thus, like many signaling proteins, CD45 can alter the signaling function depending on its location and association. To learn more about the molecular association of CD45 with the resting IgM-BCR, I had a look at the evolutionary highly conserved TMD sequence of CD45 and found that one side of the CD45 TMD carries an ICOM-related sequence in which the 2 outer AAs of the 5 LZ positions are occupied by a phenylalanine and tyrosine instead of a leucine (Fig. 2). When evaluating the CD45 ICOM, it should be kept in mind that within the lipid bilayer, a phenylalanine or tyrosine can form tight complexes with either a leucine or isoleucine residue. A similar ICOM-related sequence has also been found in the TMD of human CD101, also known as immunoglobulin superfamily member 2 (IGSF2) and cadherin 26. A related sequence is also present in the TMD of two further human PTPRs namely PTPRE and PTPRA (Tab. 3).

It has been shown that CD45 and the related transmembrane phosphatase PTPRA can form dimers and that dimerization is mediated by the extracellular domain and the TMDs of these phosphatases. It is therefore likely that the ICOM of these phosphatases is involved in their dimerization. The CD45 dimer is less active and seems to be the autoinhibited form of this tyrosine phosphatase (36). Thus, dissociation and reorganization of the CD45 dimer appears to be associated with an increased enzymatic activity of CD45. Interestingly this process is regulated by the CD45-associated protein (CD45-AP or PTPRC-AP), a small membrane protein that binds to CD45 via its leucine-rich TMD (37). Closer examination of the CD45-AP TMD sequence reveals that it carries an ICOM-related LZ structure (Tab. 3). T cell studies have shown that the binding of CD45-AP to CD45 not only leads to the dissociation of the CD45 dimer, but also to an increased phosphatase activity and CD45/TCR interaction (38). In this respect, the ICOM-mediated CD45/CD45-AP interaction is similar to that of the EpoR/gp55 pair. Thus, on resting B cells, CD45-AP may promote CD45:CD45 dissociation and the binding of CD45 to CD79b. In this way, an active tyrosine phosphatase is coupled to the IgM-BCR and this association may help to maintain the resting state of B lymphocytes.

### Model of an ICOM-Based Dynamic Regulation of B Cell Maintenance and Activation

The discovery of conserved ICOMs within the TMD of one of the two BCR signaling components and several BCR co-receptors suggests that B cell maintenance and activation is a very dynamic process involving different ICOM interactions. Based on this new information, one could imagine the following scenario for the regulation and activation of B cells via specific TMD:TMD interactions (Fig. 3). On the surface of resting B cells, the IgM-BCR may form several inhibitory complexes that prevent deregulated signaling. One complex is an autoinhibitory IgM-BCR dimer that is maintained by an ICOM-dependent CD79b/CD79b homodimerization. Within the closed dimer, the CD79b ICOMs are not accessible for interaction with CD19 or other positive signaling components. The IgM-BCR and IgD-BCR dimer could also become part of a higher oligomeric BCR structure via the interaction of the free exposed TMDs of the HC:HC’ homodimer (39). A second inhibitory complex could be formed via the ICOM-dependent interaction of the monomeric IgM-BCR with the CD45/CD45-AP complex as discussed above. Once an ICOM:ICOM negative regulatory complex is formed and delivered to the plasma membrane, it could be further stabilized by extracellular cross-linkers. A likely candidate for this function is galectin-9 (Gal9), a dimeric adaptor protein carrying two carbohydrate recognition domains separated by a flexible linker. Exposure of B cells to Gal9 suppresses IgM-BCR mobility and signaling (40). Furthermore, Gal9-deficient B cells are hyperactive, thus demonstrating the negative regulatory role of this lectin. Interestingly, several of the identified binding targets of Gal9 such as CD45, CD79b and CD180 also carry an ICOM with in their TMD. Upon B cell activation these ICOM complexes may be remodeled as outlined in Figure 3. In summary, the discovery of conserved ICOM sequences in the TMD of CD79b and several coreceptor modules suggests that the IgM-BCR regulation and activation is a dynamic process involving stable or transient ICOM-based homo- and hetero-dimerization of the IgM-BCR with itself and with CD45 and CD19.

**Figure 3.**
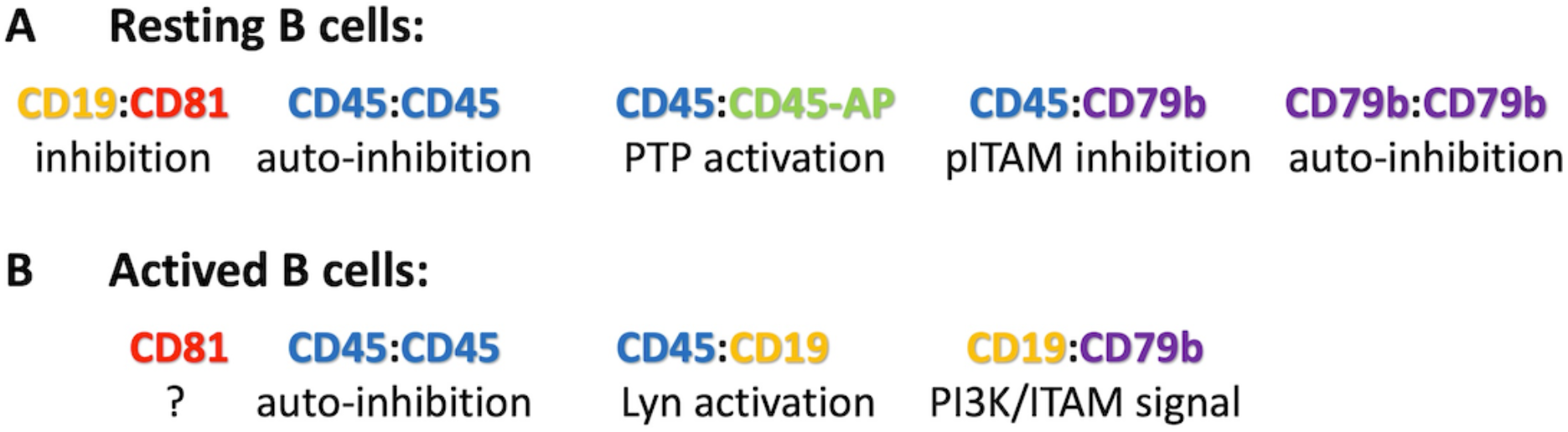
Suggested ICOM-based TMD interactions of membrane proteins on (A) resting and (B) activated B cells. A possible regulatory or signaling function of the depicted ICOM:ICOM complex is indicated below each suggested interaction.

## Discussion

Previously, it was thought that the TMDs only act as anchors for a membrane protein in the lipid bilayer. However, often a TMD sequence of a given membrane protein is evolutionary highly conserved, and this conservation suggests other important functions of a TMD besides anchoring. This has been impressively demonstrated by the recently published cryo-EM structures of the TCR and BCR complexes which revealed that the TMDs of the different receptor components occupy a considerable space and that specific TMD AA interactions play an important role in the stability and organization of the antigen receptor structure. Unfortunately, for most receptors, such as the EpoR or PDGFR, structural information is available only for the ectodomains but not for the TMD part, and this lack of information has resulted in misinterpretations of the receptor organization and function (27). For example, it was thought that most cell surface receptors are expressed as monomers that dimerize only upon ligand binding. This idea is no longer supported by more recent receptor studies. Many biochemical studies show that the TMD of single-spanning receptors form ligand-independent dimers or higher complexes in the lipid bilayer (22, 41). So far, two different TMD dimerization motifs have been studied, namely the GxxxG motif and the LZ motif (42). The GxxxG motif has been studied mainly in the context of the glycophorin A function (43, 44), whereas the LZ motif has been studied in the context of the EPO-R (45) and PDGFR (33). Interestingly, some of the ICOM-positive TMD sequences also contain a GxxxG motif and these TMDs may switch between different conformations. Compared to the GxxxG motif, the LZ motif is more flexible and can tolerate multiple AA exchanges. In particular, aliphatic and sulphur-containing AAs, such as methionine and cysteine, can increase the affinity of a TMD:TMD interaction (46, 47). This could be one of the reasons why many ICOM-positive membrane proteins on the T cell surface carry a cysteine at the 5^th^ position of the LZ structure.

The analysis described focuses on the TMD sequence of single-pass membrane proteins, but the TMDs of multi-pass membrane proteins may also carry an ICOM. However, without a detailed structural information it is not possible to determine whether the ICOMs connect the TMDs within the multi-pass receptor or establish a bridge to lateral interactors. An important class of multi-spanning membrane proteins are tetraspanins, which play an important role in the nanoscale organization of receptor clusters (48-50). Indeed, several members of this receptor family seem to carry an ICOM within their TMD (unpublished observation).

The recently determined cryo-EM structures of the monomeric TCR and BCR complexes were an important advance but they did not reveal the regulation and activation mechanism of these antigen receptors. The ICOM discovery suggests that CD79b is the main lateral interactors of the BCR. As outlined in Fig. 3, these ICOM-positive receptor components could connect the BCR to negative or positive signaling coreceptors. In the TCR complex I also identified a component that can connect this antigen receptor to multiple ICOM-positive lateral interactors (data in preparation). I propose that antigen receptors do not function as isolated entities but rather as part of an ICOM-based interactome. My hope is that further biochemical and genetic studies will confirm the role of the ICOM interactome and provide a solid foundation for a better understanding of adaptive immunity.

## Materials and Methods

Human UniProt membrane protein sequences were downloaded (https://www.uniprot.org) and filtered for unique single-pass transmembrane proteins. The TMD sequence with 5 flanking N- and C-terminal AAs were extracted using the feature annotations for each gene. This resulted in 2417 unique single-pass TMD sequences, which were scanned for ICOM motifs. Motifs were formulated as regular expressions and matched using grep.

## Acknowledgements

I thank Dr. Ying Dong for her help with the interpretation of the cryo-EM data and Dr. Peter Nielsen for his help with the ICOM motif searches. Funding was provided by the German Research Foundation (DFG) through the TRR 130 project P02 and by an RO1 grant of the National Institutes of Health (NIH) under the award number A031503.

